# Brain endothelial CXCL12 attracts protective Natural Killer cells during ischemic stroke

**DOI:** 10.1101/2021.02.18.431426

**Authors:** Shuaiwei Wang, Lauriane de Fabritus, Praveen Ashok Kumar, Yves Werner, Carole Siret, Milesa Simic, Yann M. Kerdiles, Ralf Stumm, Serge A. van de Pavert

## Abstract

Innate lymphoid cells (ILCs) have a function in homeostasis and immune responses, but their role in ischemic brain insult is unknown. Here, we report that ILCs are not resident within the mouse brain parenchyma during steady-state conditions, but are attracted after ischemic stroke. Specifically, we identified NK cells, ILC1s, ILC2s and ILC3s within the lesion, the highest influx being observed for NK cells and ILC1s. We further show that Cxcl12 expressed at the blood-brain barrier is essential for NK cells and NKp46^+^ ILC3s to migrate toward the lesion. Complementary, *Cxcr4*-deficiency in NK cells prevented NK cells from entering the infarct area. The lack of NK cell migration resulted in a higher neurological deficit in the beam-walk sensorimotor test. Our data show a new role for blood-brain barrier-derived Cxcl12 in attracting protective NK cells to ischemic brain lesions and identifies a new Cxcl12/ Cxcr4-mediated component of the innate immune response to stroke.

## Introduction

Ischemic stroke triggers prominent innate immune responses that plays multiphasic roles in the pathogenesis of ischemic brain injury. Microglia are rapidly activated at the early stage after stroke and the activation persists at the later phase, resulting in dual effects for the injury (Eldahshan et al., 2019). Subsequently, leukocyte including neutrophils, monocytes and dendritic cells (Garcia et al., 1994; Gelderblom et al., 2009; Eldahshan et al., 2019; Werner et al., 2020) are attracted to the lesioned tissue. While neutrophil recruitment is thought to be deleterious, brain infiltration by anti-inflammatory monocytes has been connected to improved outcomes after experimental stroke. Whether besides NK cells, any other ILC was present during stroke is not known. Since they are critical for the maintenance of homeostasis and tissue remodeling as well as tissue repair under certain circumstances (Vivier et al., 2018), they can either be beneficial or detrimental to the outcome of the experimental stroke. Recently, NK cells have been reported in neurological diseases, including experimental autoimmune encephalomyelitis (EAE), Alzheimer and Amyotrophic Lateral Sclerosis (ALS) and stroke (Gan et al., 2014; Romero-Suárez et al., 2019; Gate et al., 2020; Garofalo et al., 2020). ILC1s and ILC3s are also involved in EAE (Hatfield and Brown, 2015; Romero-Suárez et al., 2019) and ILC3s were shown to be critical for the Th17 response in EAE (Kwong et al., 2017). However, the mechanism involved in recruitment of ILCs within the infarct region remains largely unknown. The expression chemokine CXCL12 and its receptor CXCR4 within the brain increased after ischemic stroke onset, which plays a critical role in the migration of monocytes to brain parenchyma (Werner et al., 2020). CXCL12 was shown to be expressed by the blood brain barrier (BBB) endothelial cells at steady state and redistributed in location during EAE in mice or multiple sclerosis in humans(Williams et al., 2014), however a role for BBB derived CXCL12 during ischemic stroke was not known. Given that CXCR4 is expressed by ILCs, and the antagonist AMD3100 treatment affects the cell numbers of NK cells in the ischemic hemisphere (Inngjerdingen et al., 2001; Klein and Rubin, 2004; Selvaraj et al., 2017), CXCL12/CXCR4 signaling could directly regulate NK cell and ILC migration to the infarct region. Indeed, CXCR4^+^ NK cells were shown to migrate in patients specifically during the remission phase of multiple sclerosis(Serrano-Pertierra et al., 2015), and were also observed in stroke regions in patients (Gan et al., 2014). However, if they respond to Cxcl12 expressed in the blood-brain barrier after ischemic lesion is unknown. Even though ILC populations have been reported in central nervous system (CNS) disorders including EAE and Alzheimer, there is an apparent lack in knowledge on the involvement of ILCs in stroke, and if they are involved in tissue repair or the deleterious immune reaction.

In this study, we show that NK cells, ILC1s, ILC2s, NKp46^+^ ILC3s and LTi cells were recruited to stroke brain, the majority being NK cells and ILC1s. Using *Cdh5^CreERT2^;Cxcl12^flox^* mice in which *Cxcl12* was specifically deleted in BBB endothelial cells and *Cxcr4* deficient NKp46^+^ cells, we demonstrated that BBB CXCL12-CXCR4 axis was essential for the influx of NK cells into the stroke lesion. We further showed that these NK cells protect the brain and improved mouse motor movement after stroke induction.

## Results

### ILCs are present within the stroke lesion

To analyze whether ILCs are involved in responses towards ischemic stroke in a reproducible and minimally invasive manner, we used photothrombotic (PT) induced stroke. We injected Rose Bengal mice in the ischemic group whereas the mice in the sham group were only injected with saline. Subsequently, intense light with 1mm diameter illuminated a precise position on the skull above the hippocampus to photo-activate Rose Bengal (Fig. 1A). The cortical lesion was identified by Nissl staining on vibratome sections from stroke brain at post-operative day 2 (P2) (Fig. 1B). We determined the ILC presence in or near the lesion by immunofluorescence (Fig. 1C-E) and a detailed time course on the ILC influx was investigated by flow cytometry analysis (Fig. 2).

**Figure 1:**
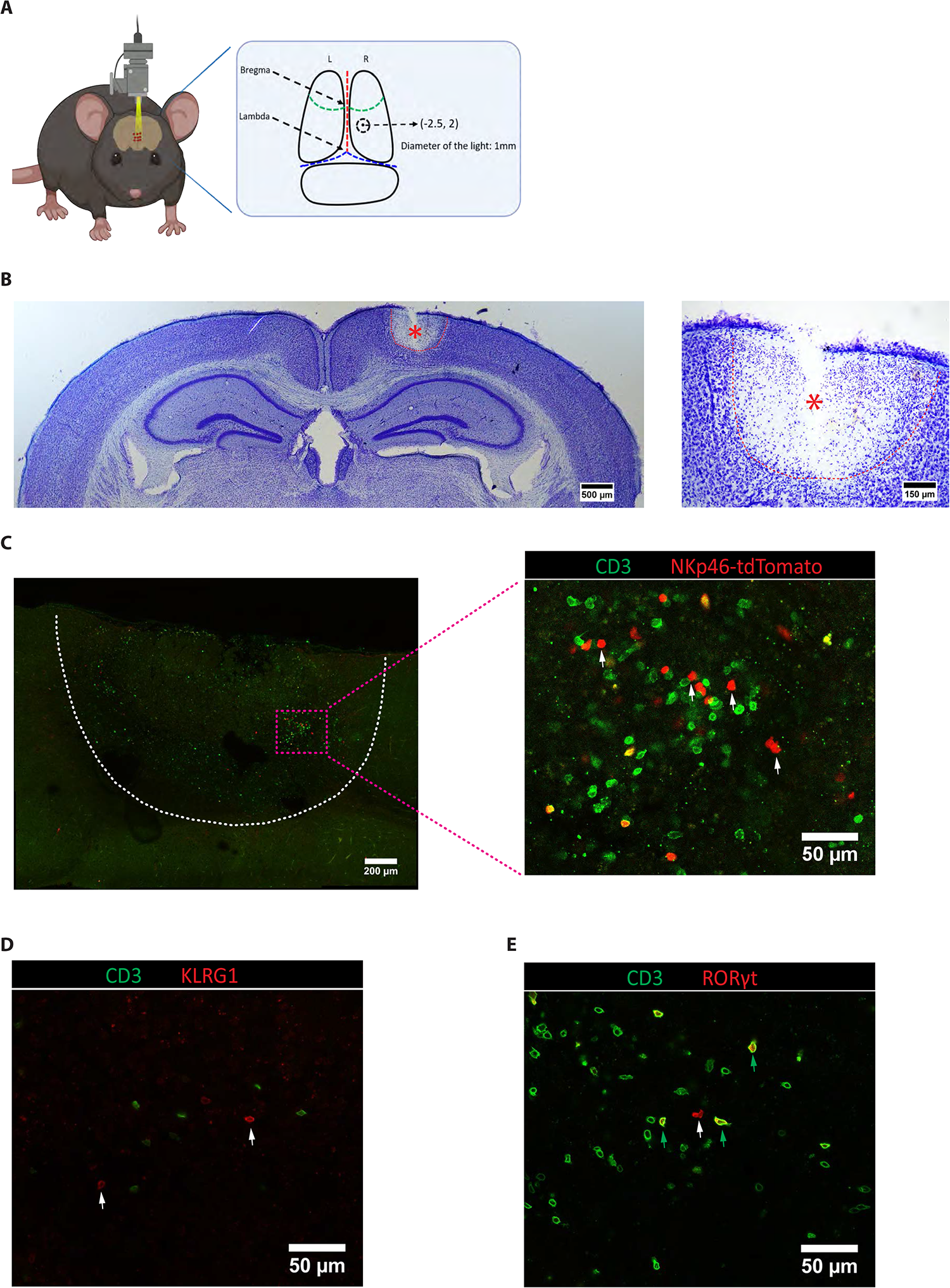
Presence of ILCs in the ischemic stroke lesion. (A) Coordinates of the lesion on the brain of photothromobotic (PT) mouse model at 2.5mm posterior to bregma and 2.0mm lateral to midline with 1mm diameter. (B) Nissl staining on the vibratome section to indicate the lesion in the stroke brain at P2. (C) Immunofluorescence on the vibratome sections of stroke brain at P10. Since the markers of ILCs, NKp46, KLRG1 and RORγt are also expressed by CD3^+^ non-ILCs, we used CD3 to distinguish NK/ILC1s from NKT cells, ILC2s from regulatory T cells (Tregs) and NKp46^+^ ILC3s from Th17 cells. CD3^+^NKp46^+^ cells (NKT, green arrows) and CD3^−^NKp46^+^ cells (NK, ILC1 or NKP46^+^ILC3, white arrows), (D) CD3^−^KLRG1^+^ cells (ILC2, white arrows), (E) CD3^+^RORγt^+^ cells (Th17, green arrows) and CD3^−^RORγt^+^ cells (ILC3, white arrows). Data shown are from at least two independent experiments.

**Figure 2:**
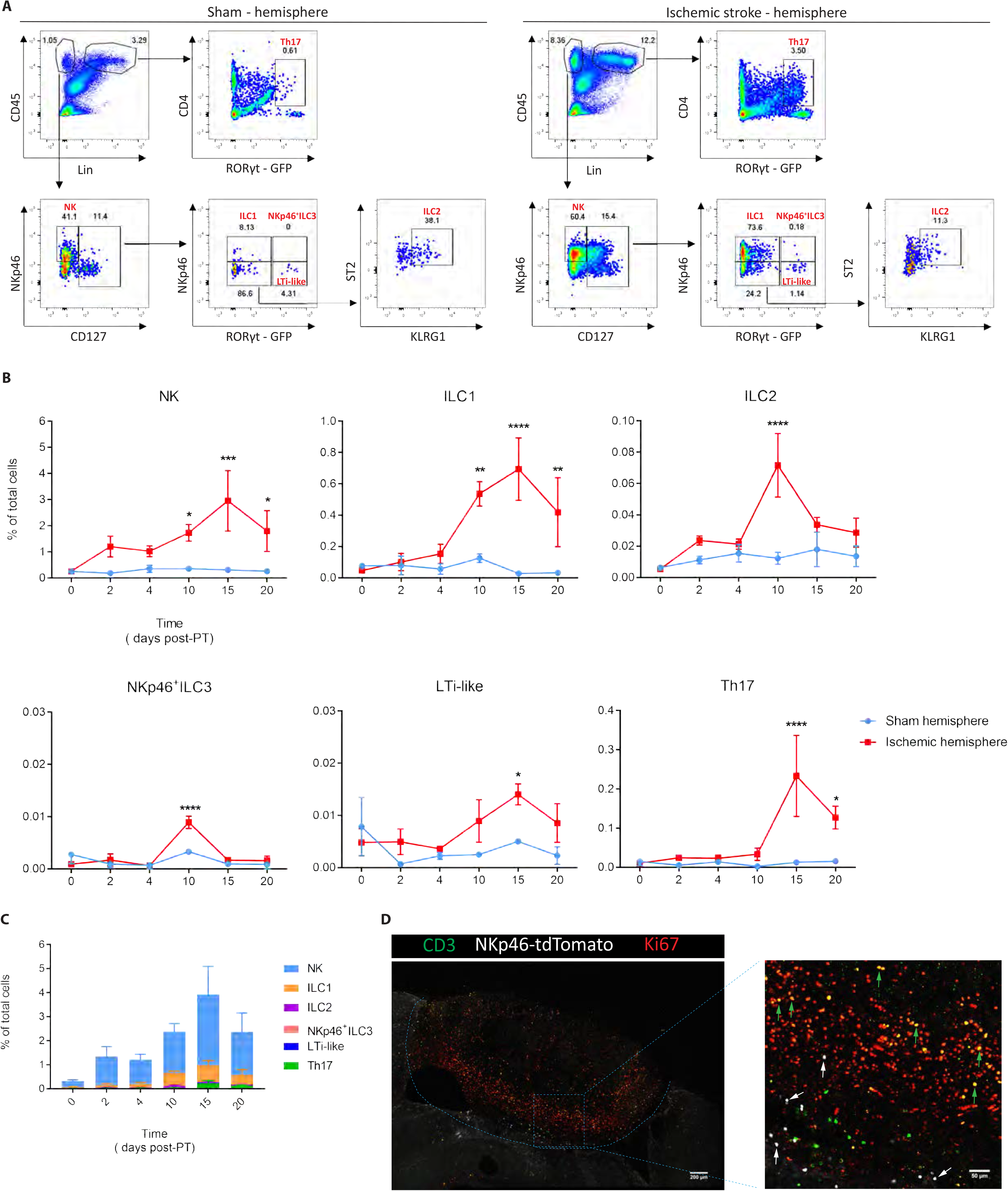
Time-course of ILC migration towards the ischemic stroke lesion. Cytometry analysis of ILCs in the sham (A, left) and P15 stroke hemisphere (A, right), NK (CD45^+^Lin^−^NKp46^+^CD127^−^), ILC1 (CD45^+^Lin^−^NKp46^+^CD127^+^RORγt^−^), ILC2 (CD45^+^Lin^−^NKp46^−^CD127^+^RORγt^−^ST2^+^KLRG1^+^), NKp46^+^ILC3 (CD45^+^Lin^−^NKp46^+^CD127^+^RORγt^+^), LTi-like (CD45^+^Lin^−^NKp46^−^CD127^+^RORγt^+^), and Th17 (CD45^+^Lin^+^CD4^+^RORγt^+^). Representative dot plots of ILCs and Th17 cells on P15 are shown with the averages for the gates. (B) Quantification of ILC populations and Th17 cells in the hemisphere after PT induction. Data represent n=3 mice. NK: P10 *P=0.0364, P15 ***P=0.0003, P20 *P=0.021; ILC1: P10 **P=0.0052, P15 ****P<0.0001, P20 **P=0.008; ILC2: P10 ****P<0.0001; NKp46^+^ILC3: ***P=0.0001; LTi-like: P15 *P=0.0232; Th17: P10 ****P<0.0001. (C) Comparison of ILC and Th17 cell numbers in stroke hemisphere at different time points after induction. Data represent n=3 mice. (D) Confocal images showing immunofluorescence for CD3, NKp46 and Ki67 in the PT-induced lesion at day 10. Green arrows point to the proliferating CD3^+^ cells, while red arrows point to the non-proliferating CD3^−^NKp46^+^ cells including NK cells, ILC1s and ILC3s. Data are representative of two independent experiments.

CD3^−^NKp46^+^ NK, ILC1 and −3 cells were identified in the stroke lesion (Fig. 1C). The presence of CD3^−^KLRG1^+^ cells in the stroke lesion indicated that ILC2s were recruited to the stroke brain (Fig. 1D). The transcription factor RORγt expressed in ILC3 and Th17 cells was identified by using *RORc^eGFP^*-reporter mice. We observed CD3^−^RORγt^+^ and CD3^+^RORγt^+^ cells in the stroke lesion (Fig. 1E) and indicated that both ILC3 and Th17 cells were attracted to ischemic stroke brain. The presence of very few ILC3s and many CD3^−^NKp46^+^ cells in the stroke lesion suggests that a large proportion of CD3^−^NKp46^+^ cells are NK and/or ILC1s (Fig. 1C and 1E). To comprehensively understand the spatial distribution of ILCs in the brain lesion after ischemic stroke, we performed whole-mount immunofluorescence on the brain. 3D imaging analysis showed the accumulation of CD3^−^NKp46^+^ (NK, ILC1 and NKp46^+^ILC3), CD3^−^KLRG1^+^ (ILC2), CD3^−^RORγt^+^ (ILC3 and LTi) cells, the majority of which were located at the border of the lesion at P2 (Video S1) and P10 (Video S2).

### Time course of ILC migration to the stroke hemisphere by flow cytometry

To quantify the ILC presence at different time points within the lesioned brain, we used a flow cytometry staining panel which identified all ILC subsets (gating strategy in Fig. S1A). Using *RORc^eGFP^*-reporter mice, we analyzed CD4^+^RORγt^+^ Th17 cells to assess the validity of our PT stroke model. NK cells, ILC1s, ILC2s, NKp46^+^ ILC3s and NKp46^−^ LTi-like cells were analyzed (Fig. 2A). At P0, we observed no specific ILC migration towards the brain and general ILC numbers were either non-existent or very low (NK cells). NK cell numbers slightly increased at P2 (Fig. 2B) and peaked at P15 (Fig. 2A and 2B). ILC1 numbers increased at P10 and peaked at P15 (Fig. 2B) but were not detected at P2 and P4 (Fig. 2B). ILC2 numbers sharply increased after P4 and lowered at P15. The number of NKp46^+^ ILC3 cells peaked at P10 while NKp46^−^ LTi-like cells peaked at P15 (Fig. 2B). Th17 cell numbers significantly increased at P15 and decreased at P20 (Fig. 2B). The presence of Th17 cells was consistent with previous data using the middle cerebral artery occlusion (MCAO) stroke model (Fig. 2B)(Ito et al., 2019). Cranial dura mater is the outmost layer of cranial meninges, which contains the dural venous sinuses draining blood and shunted lymphatic vessels draining cerebrospinal fluid (CSF) from brain (Aspelund et al., 2015; Louveau et al., 2015; Da Mesquita et al., 2018). As there is increasing evidence for the role for dural meninges in immunity (Alves De Lima et al., 2020), we quantified ILCs in the dura mater after PT induction. We did not observe a significant ILCs increase nor Th17 cells in the ischemic dura mater (Fig. S1B, S2A andS2B). Immune cells are drained from the brain to the deep cervical lymph nodes (dcLN) and mandibular lymph nodes (mandiLN) (Ma et al., 2017). An increase in ILC numbers within these LN indicate an active clearance and/or migration of the ILCs. However, ILC and Th17 cell numbers were not increased within dcLN or mandiLN draining the ischemic part of the brain (Fig. S1C, S2C-E), similar as mesenteric (mLN) and axillary lymph nodes (axiLN) (Fig. S2F and S2G). Summarized, analysis of ILC subsets in the stroke hemisphere at different time points after PT stroke showed that NK cells and ILC1s constituted the majority of the ILCs responding to the stroke lesion in the brain (Fig. 2C). Ki67 immunofluorescence staining showed that some CD3^+^ cells were proliferating, but CD3^−^NKp46^+^ cells were not (Fig. 2D). Our data suggests that the progressive increase of NKp46^+^ ILCs is caused by migration towards the lesion rather than *in situ* proliferation.

### The role of endothelial CXCL12 in migration of NK cells and NKp46^+^ ILC3s

Using *in situ* hybridization *Cxcl12* was observed throughout the brain and increased after stroke (Werner et al., 2020). Using *Cxcl12-DsRed* reporter mice, we observed CXCL12 on CD31^+^ endothelial cells within the stroke lesion at P10 (Fig. 3A). This indicated that CXCL12 is generated by the BBB to attract lymphocytes, as was shown before in EAE and MS(Klein and Rubin, 2004). To study whether the migration of NKp46^+^ ILCs was mediated by BBB derived CXCL12, we specifically and conditionally deleted endothelial *Cxcl12* using the *Cdh5^CreERT2^;Cxcl12^flox/flox^* model (from now named *Cxcl12^Cdh5−/−^*) since the full *Cxcl12* knock-out is embryonic lethal. 4-Hydroxytamoxifen (4OHT) was injected 3 weeks before PT induction (Fig. 3B) so that the 4OHT was washed out before the experiment. To confirm *Cxcl12* deletion, we sorted CD45^−^CD31^+^ endothelial cells (Fig. S3A and S3B) and performed a RT-qPCR analysis which confirmed lack of *Cxcl12* within the endothelial cells (Fig. 3C). NK cells, intILC1s, ILC1s, ILC2s and NKp46^+^ ILC3s were analyzed by flow cytometry (Fig. 3D and S4A). Compared to the controls, we observed a decrease of NK cells in the *Cxcl12^Cdh5−/−^* stroke brain at P2 and a significant decrease at P15 (Fig. 3E), indicating that the endothelial-produced CXCL12 is directly involved in the attraction of NK cells. A significant decrease of NK cells in the dcLN, mandiLN and mLN was not observed (Fig. S5D-F). NK cells in the dura mater and axiLN (Fig. S5A and S5C) significantly decreased at P2 (Fig. S5B and S5G). No difference on intILC1 and ILC1 migration was detected at P2 and P15 (Fig. 3E), suggesting that these two subsets were attracted to the stroke brain through other mechanisms. IntILC1s were decreased in the dura mater and axiLN at P2 (Fig. S5B and S5G). ILC1s were also decreased in the dura mater and mandiLN (Fig. S5B and S5E). ILC2 migration was not affected in the *Cxcl12^Cdh5−/−^* stroke brain, dura mater and lymph nodes (Fig. 3E, S5B and S5D-G). NKp46^+^ ILC3s were decreased within stroke brain at P15 and axiLN at P2 (Fig. 3E and S5G).

**Figure 3:**
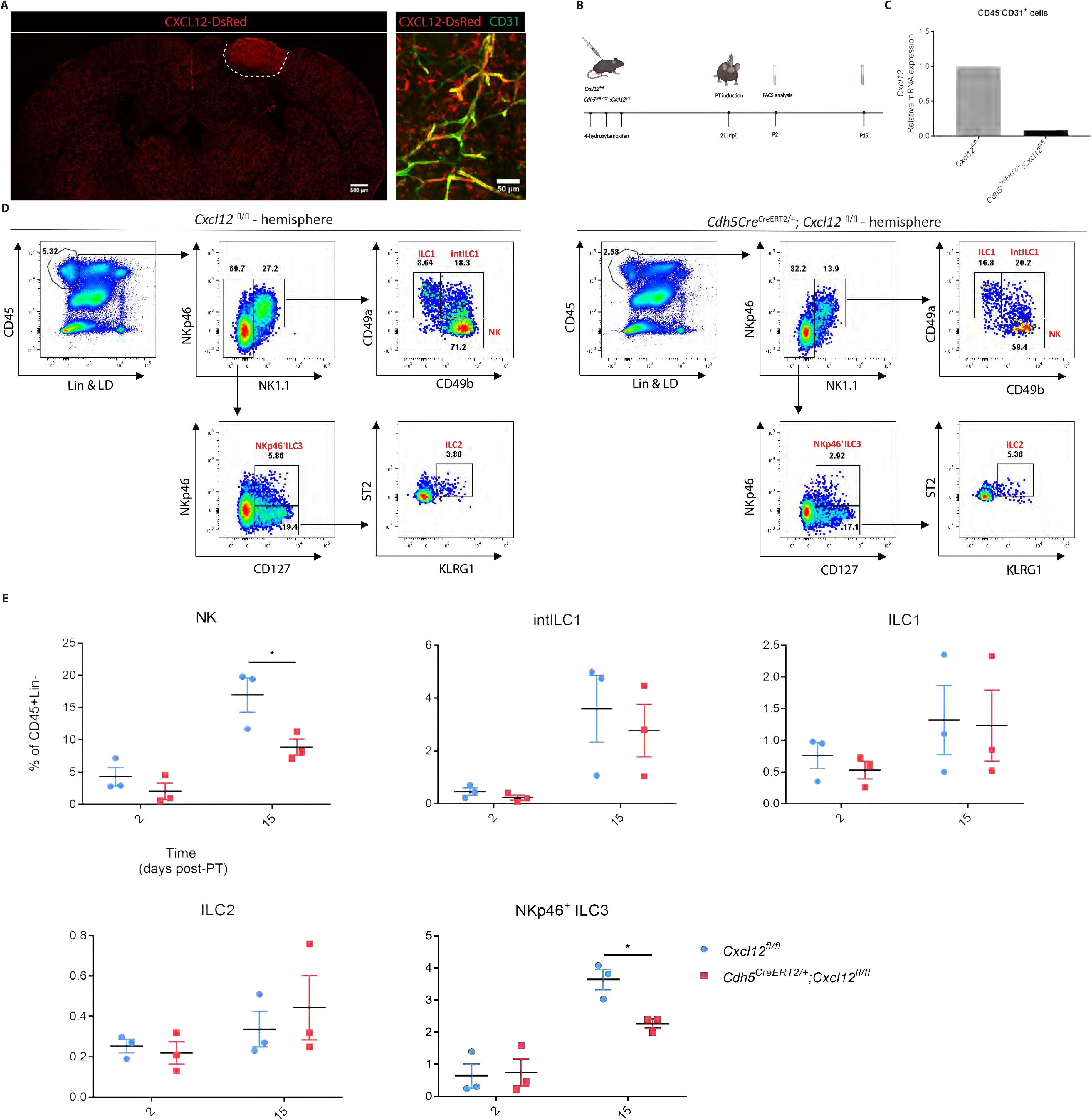
Recruitment of ILCs via blood brain barrier expressed CXCL12. (A) CXCL12 expression within the lesion at P10, co-localizing with CD31^+^ endothelial cells in *Cxcl12^DsRed^* mouse. (B) Cxcl12^fl/fl^ and *Cdh5^CreERT2/+^;Cxcl12^fl/fl^* mice were injected with 4OHT at day −21 relative to PT induction and analyzed by flow cytometry at day 2 and 15 after PT induction. dpi: days post injection (C) *Cxcl12* mRNA levels in the sorted endothelial cells from WT (Cxcl12^fl/fl^) and KO (*Cxcl12^Cdh5–/–^*) mice. RNA was extracted from cells obtained from 3 mice. (D) Flow cytometry analysis of ILCs in the P15 stroke brain from WT (*Cxcl12^fl/fl^*) and KO (*Cxcl12^Cdh5–/–^*) mice, NK (CD45^+^Lin^−^NKp46^+^NK1.1^+^CD49a^−^CD49b^+^), intILC1 (CD45^+^Lin^−^NKp46^+^NK1.1^+^CD49a^+^CD49b^+^), ILC1 (CD45^+^Lin^−^NKp46^+^NK1.1^+^CD49a^+^CD49b^−^), ILC2 (CD45^+^Lin^−^NKp46^−^NK1.1^−^CD127^+^ST2^+^KLRG1^+^) and NKp46^+^ILC3 (CD45^+^Lin^−^NK1.1^−^NKp46^+^CD127^+^). (E) Quantification of ILCs in the stroke hemisphere from Control (*Cxcl12^fl/fl^*) and KO (*Cxcl12^Cdh5−/−^*) mice, at day 2 and 15 after PT induction. Each data point represents an individual mouse. NK *P=0.0113, NKp46^+^ ILC3 *P=0.0186.

### Migration of NK cells to the stroke lesion depends on CXCR4

Of the ILC family, mainly NKp46^+^ NK cells and ILC1s responded to the stroke lesion (Fig. 2C). Therefore, we used the *Ncr1^iCre^* model to delete *Cxcr4* specifically in these cells. We confirmed *Ncr1^iCre^* validity with the *Ncr1^iCre/+^; Rosa^tdTomato/+^* reporter model. At P10, 96.4 ± 0.4% of NK cells were tdT^+^ (Fig. 4A). Of the total CD45^+^ population, more than 95% of tdT^+^ cells were lineage negative (Fig. 4B), of which 86.9 ± 2.5% constituted NK cells as well as ILC1s and 7.0 ± 1.5% constituted NKp46^+^ ILC3s(Fig. 4B). Therefore, we concluded that the *Ncr1^iCre/+^* mice is a suitable model to drive the specific depletion of *Cxcr4* in almost all the NKp46^+^ ILCs. We confirmed the lack of *Cxcr4* expression by sorting the CD45^+^Lin^−^NKp46^+^ cells (Fig. S6A and S6B) from *Ncr1^iCre/+^;Cxcr4^flox/flox^*, hereafter named *Cxcr4^Ncr1−/−^* mice, and observed drastically reduced *Cxcr4* expression by RT-qPCR (Fig. 4C). Although the fluorescence of cell surface NKp46 was lower in the *Ncr1^iCre/+^* mice, NK and ILC1 cell numbers in the *Ncr1^iCre/+^* mice were comparable with *Cxcr4^flox/flox^* mice (Fig. S7A) as reported before in the original description of this line (Narni-Mancinelli et al., 2011). We induced PT on control (*Cxcr4^flox/flox^* and *Ncr1^iCre/+^*) and *Cxcr4^Ncr1−/−^* mice and analyzed the ILC subsets by flow cytometry (Fig. S4A). We observed that deletion of *Cxcr4* resulted in loss of NK cell migration, while in the control mouse brains NK cell numbers increased at P2 and P15 after PT induction (Fig. 4D, 4E and S7B). We also observed a decrease in NK cell numbers in the dura mater at P2 and P15 and in the draining dcLN at day 2 (Fig. S8A-D). At P15 we detected fewer intILC1s in the *Cxcr4^Ncr1−/−^* brain (Fig. 4E) although no significant difference in the intILC1s numbers were observed in the dura mater, dcLN, mandiLN, mLN and axiLN (Fig. S8A-G). ILC1s, ILC2s and NKp46^+^ ILC3s were not changed in the brain, dura mater, dcLN, mandiLN, mLN and axiLN of *Cxcr4^Ncr1−/−^* mice (Fig. 4E and S8A-G).

**Figure 4:**
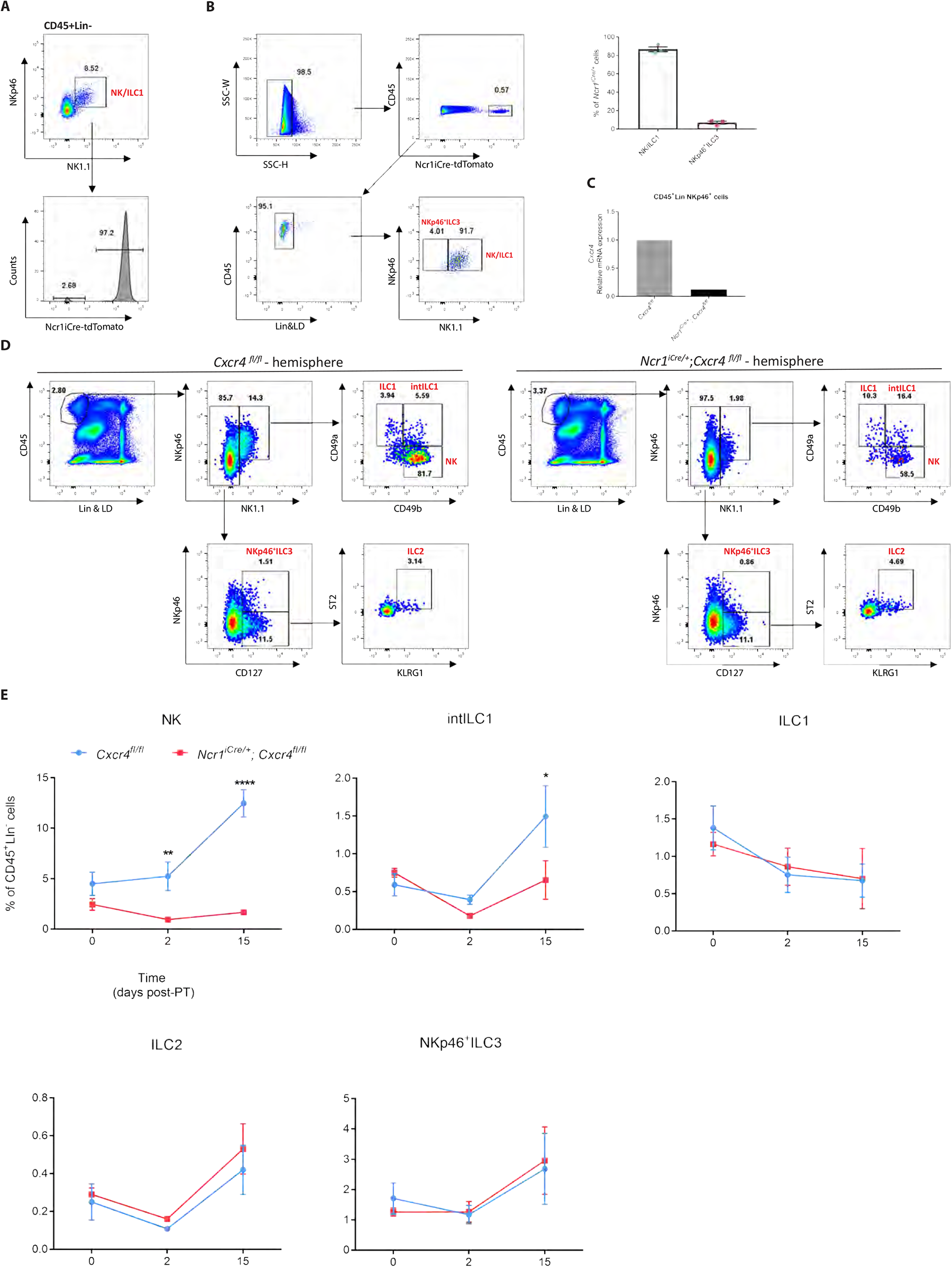
NK cell migration is CXCR4-dependent. (A) Testing the *Ncr^iCre^* efficiency in the NKp46^+^ cells by FACS using the *Rosa^tdTomato^* reporter. (B) Analysis of the NKp46^+^ cell populations in the *Ncr^iCre^*; *Rosa^tdTomato^* reporter. Representative dot plots at P15 are shown. Each data point represents an individual mouse. (C) *Cxcr4* mRNA levels in sorted NK cells from Control (*Cxcr4^fl/fl^*) and KO (*Cxcr4^Ncr1−/−^*) mice. RNA was extracted from cells obtained from 3 mice. (D) Flow cytometry analysis of ILCs in the stroke hemisphere from Control (*Cxcr4^fl/fl^*) and KO (*Cxcr^4Ncr1−/−^*) mice at P15, NK (CD45^+^Lin^−^NKp46^+^NK1.1^+^CD49a^−^CD49b^+^), intILC1 (CD45^+^Lin^−^NKp46^+^NK1.1^+^CD49a^+^CD49b^+^), ILC1 (CD45^+^Lin^−^NKp46^+^NK1.1^+^CD49a^+^CD49b^−^), ILC2 (CD45^+^Lin^−^NKp46^−^NK1.1^−^CD127^+^ST2^+^KLRG1^+^) and NKp46^+^ ILC3 (CD45^+^Lin^−^NK1.1^−^ NKp46^+^CD127^+^). Representative dot plots of ILCs at P15 are shown. (E) Quantification of ILCs in the stroke hemisphere from WT (*Cxcr4^fl/fl^*) and KO (*Cxcr4^Ncr1−/−^*) mice at P0, P2 and P15. Data represent n=3 mice. NK P2 **P=0.0085, P15 ****P<0.0001.

### NK cells positively regulate behavioral deficits after PT stroke induction

To establish the role of NK cell presence on behavior during ischemic stroke, we performed a beam-walk sensorimotor test. We induced ischemic stroke on the entire cortical area which controls forelimb and hindlimb movement by using a 2mm diameter light-beam window in Rose Bengal injected mice (Fig. 5A and 5B). We did not observe a difference in ILC numbers between this model and the previously used 1mm diameter window for ischemic stroke (data not shown). The mice used in this study were trained for 4 consecutive days before PT induction and the analysis of their motor behavior during the beam-walk sensorimotor test was assessed by video and compared to that just before PT induction (day 0). We recorded the videos every second day and counted the total steps as well as every faulty step or slip in each test. We observed that loss of NK cells in the *Cxcr4^Ncr1−/−^* mice negatively affected behavioral deficits after PT induction (Fig. 5C), indicative of a positive influence of NK cells during on ischemic stroke. In order to confirm the effect of NK cells on motor behavior and to confirm that the effect observed was not due to a possible effect of the *Cxcr4^Ncr1−/−^* line on NK cell hematopoiesis and NK cell numbers in general, we depleted NK cells in C57BL/6J mice by anti-NK1.1 treatment 1 week and 1 day before PT induction. Loss of NK cells was observed before and throughout the beam-walk sensorimotor test analysis (Fig. 5D, 5E and S9). The mice injected with anti-Nk1.1 and concomitantly loss of NK cells showed more severe behavioral deficits at P2 (Fig. 5F), confirming the protective effect of NK cells.

**Figure 5:**
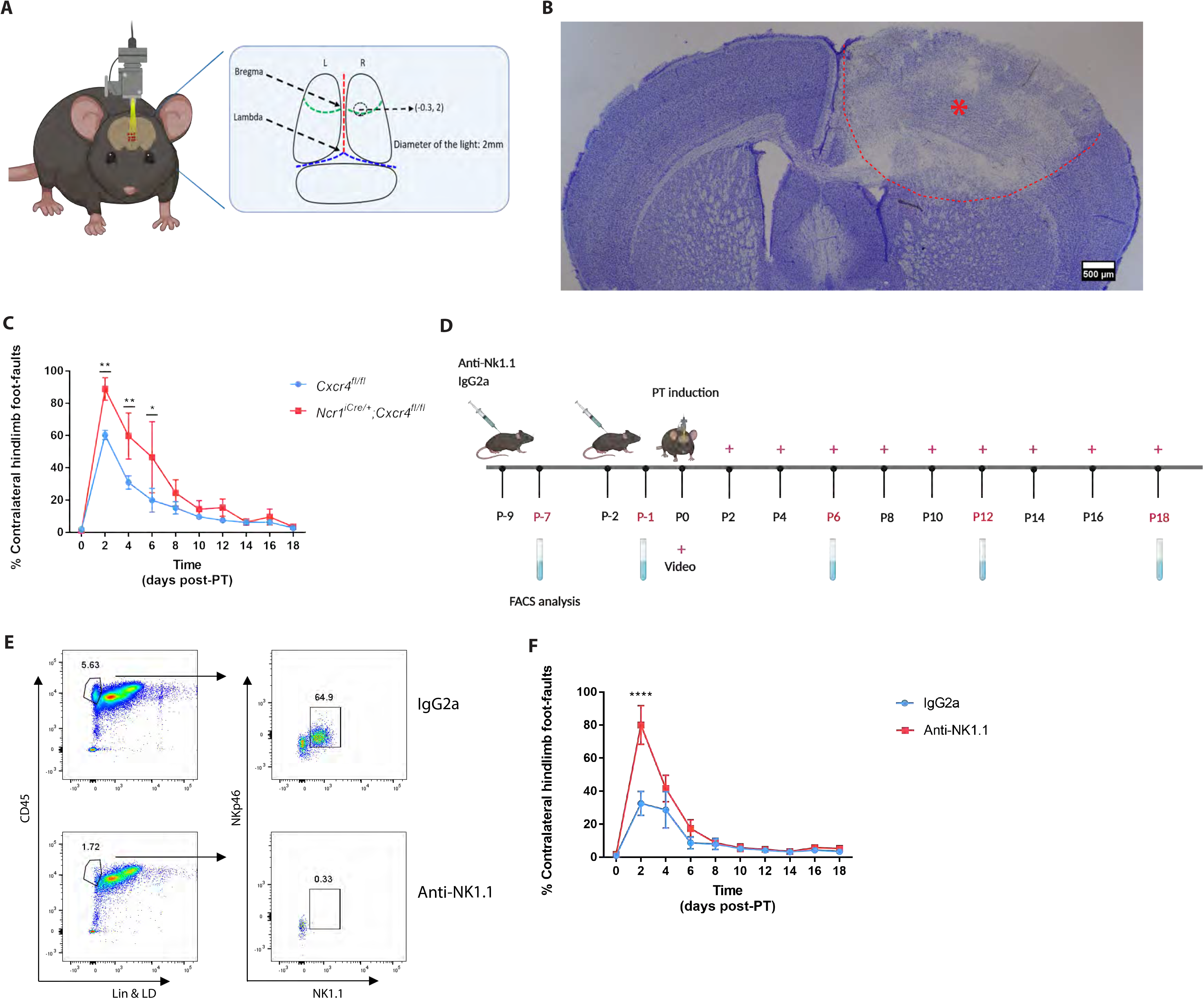
NK cells positively regulate behavioral deficits after PT stroke induction. (A) Coordinates of the lesion on the brain of photothromobotic (PT) mouse model used for beam-walk sensorimotor test analysis. The center of the lesion is 0.3mm in front of bregma and 2.0mm lateral to midline, with the 2mm of diameter. (B) Nissl staining on the vibratome section of stroke brain at day 2 after PT. (C) Beam-walk sensorimotor test analysis of Control and KO mice after PT induction by calculating the percentage of contralateral hindlimb faults. Data represent n=4 mice. P2 **p=0.0058, P4 **P=0.0053, P6 **P=0.0096. (D) Anti-NK1.1 or IgG2a was injected at day 9 (P-9) and day 2 (P-2) before PT induction and the efficiency of depletion in blood (E) was tested at P-7, P-1, P6, P12 and P18. The videos were recorded just before PT induction (P0) and every second day afterwards. (F) The beam-walk sensorimotor test analysis was performed by calculating percentage of contralateral hindlimb faults. Data represent n=5 mice. P2 ****P<0.0001.

## Discussion

Innate immune cells like neutrophils and macrophages are associated with an immune reaction after induction of ischemic stroke in the brain (Werner et al., 2020). However, the presence and role of ILCs in ischemic stroke brain was unknown. In this study, we thoroughly analyzed the presence of all ILCs in experimental ischemic stroke within the brain and determined the kinetics of ILCs migration towards the stroke lesion. We have used a minimally invasive and reproducible mouse model using photo-thrombosis (PT), which mainly affected the cortex. In this model, we observed the presence of NK/ILC1s, ILC2s and ILC3s within the brain lesion.

ILCs are considered tissue resident (Gasteiger et al., 2015), although some circulation was reported (Dutton et al., 2019). Conversely, we observed very few NK cells and hardly any ILCs in the brain at P0, or in naïve conditions. Therefore, ILCs observed in the brains were not resident and migrated towards the lesion after the stroke induction. We observed that NK cells and ILC1s were the main ILC populations increased after PT stroke, peaking at P15 and outnumbering the Th17 population. The increase in NK and Th17 cells was consistent with the increase found in the tMCAO model for ischemic stroke (Gan et al., 2014; Liu et al., 2017; Ito et al., 2019), confirming our PT model for analyzing ILCs during ischemic stroke in the brain. Even though ILC2s, NKp46^+^ ILC3s and LTi-like cells also increased, they represented a small portion of the infiltrated ILCs. The lack of Ki67 suggest that the robust accumulation of NK cells and ILC1s in the stroke brain was not caused by in situ proliferation, but rather progressive migration towards the ischemic lesion. This is consistent with a previous study in which a correlation between spleen atrophy and presence of NK cells after ischemic stroke in the brain was observed (Liu et al., 2017), indicating mobilization from the spleen towards the lesion. In the EAE model it was shown that lymphocytes accumulated within the dura near the lesion (Kwong et al., 2017). Conversely, in our PT model we did not observe an increase of ILCs within the dura during ischemic stroke.

CXCL12 was reported to be involved in the migration of CXCR4^+^ monocytes towards stroke (Werner et al., 2020). Moreover, in EAE it was shown that CXCL12 from the BBB was involved in the migration of the lymphocytes towards the lesion(Klein et al., 2017), but the role for CXCL12 from the BBB in the attraction of innate immune cells was not shown before. We deleted CXCL12 in the BBB by using *Cxcl12^Cdh5−/−^* mice, which resulted in decreased NK and NKp46^+^ ILC3 numbers. Even though ILC1s express CXCR4, their migration was not affected in *Cxcl12^Cdh5−/−^* nor in *Cxcr4^Ncr1−/−^* mice. This suggests that their migration to the ischemic lesion occurred through other mechanisms, possibly CX3CL1/CX3CR1 signaling as was shown before for NK cells and intILC1s (Gan et al., 2014). The intILC1s were decreased in the ischemic lesions in *Cxcr4^Ncr1−/−^* mice, while not affected in *Cxcl12^Cdh5−/−^* mice. Possibly, intILC1s require a different CXCL12 source than expressed by the BBB, as it is expressed throughout the brain (Stumm et al., 2002). The lack of NK cells in *Cxcr4^Ncr1−/−^* mice to the lesion in this model suggest that the BBB is essential for the migration of NK cells to the stroke lesion by CXCL12 expression.

The role for NK cells became clear when we prevented migration of NK cells in *Cxcr4^Ncr1−/−^* and by anti-NK1.1 antibody treatment. Using these complementary models, we demonstrated a protective effect of NK cells on recovery of motor behavior *in vivo*. It was shown that NK cells induce homeostatic anti-inflammatory activities of astrocytes via IFN-γ (Hindinger et al., 2012; Sanmarco et al., 2021). Indeed, astrocytes were found to be activated in ischemic stroke already at P3 (Werner et al., 2020), at the same time when the effect of NK cell absence was observed in our beam-walk sensorimotor test. Thus, NK cells could dampen the inflammation via astrocytes in the stroke region to promote recovery of motor control. A disadvantage of the beam-walk sensorimotor test is the fast recovery in motor function of mice after stroke. Therefore, in control, *Cxcr4^Ncr1−/−^* and anti-NK1.1 depleted mice, recovery occurred before we could assess *in vivo* the functional relevance of NK cells at P15. Interestingly, a study on T-cell involvement in ischemic stroke, using the *Rag1^−/−^* model lacking all adaptive lymphocytes but which still had NK cells, showed a lower neurogenic deficiency (Kleinschnitz et al., 2010). This data confirms that NK cells in this model could positively affect recovery in this testing. Conversely, in another study it was shown that NK cells had a detrimental effect on stroke (Gan et al., 2014). In this study *Rag2^−/−^γc^−/−^* animals, which constitutively lacked all lymphocytes including NK cells, and anti-NK1.1 treatment was used and tested with the Bederson test. The beam-walk sensorimotor test was considered a reliable measurement while the Bederson test was considered more subjective (Schaar et al., 2010). The different genetic strains and behavioral test used could have contributed to this difference.

In this study, we thoroughly examined the presence of all ILCs in ischemic stroke. ILCs are not resident within the brain and during ischemic stroke, we observed mainly NK cells and, in lesser extent, intILC1s and ILC1s located at the rim of the lesion. CXCL12 expressed by the BBB attracted the NK cells directly towards the lesion. Deleting this chemokine in endothelial cells, as well as the receptor on the NKp46^+^ cells, prevented migration towards the lesion. CXCR4 is important for the attraction of the NK cells, and other innate immune cells towards stroke, indirectly affecting microglial function (Werner et al., 2020) and was shown in both studies to be positively associated with stroke recovery. NK cells have been detected in ischemic lesions in humans (Gan et al., 2014; Liu et al., 2017), and it was suggested to restrict NK cell migration towards the lesion to prevent secondary pneumonia using e.g. AMD3100 (Liu et al., 2017). However, our data indicate that the migration of CXCR4^+^ NK cells could be beneficial for recovery. The positive effects of NK cells were shown in EAE before (Lünemann and Münz, 2008). Similarly, CXCR4^+^ NK cells were observed specifically during the remission phase in multiple sclerosis (Serrano-Pertierra et al., 2015). If the role of NK cells and other ILC1 members is similar in ischemic lesions of patients remains to be established.

## Materials and methods

### Mice

*RORc^eGFP^* mice were kindly provided by Gérard Eberl (Pasteur Institute, Paris, France)(Sparwasser et al., 2004). *Ncr1^iCre^* mice were kindly provided by Eric Vivier (CIML, Marseille, France)(Narni-Mancinelli et al., 2011). *Cdh5^CreERT2^* mice were kindly provided by Ralf Adams (Max Planck Institute for Molecular Biomedicine, Münster, Germany)(Wang et al., 2010). *Cxcr4^flox^*, *Cxcl12^flox^* and *Rosa^tdTomato^* mice were obtained from Jackson Laboratory(Nie et al., 2004; Greenbaum et al., 2013; Madisen et al., 2010). Except for the 3D imaging analysis, 8-14 weeks old adult male mice were used. All the mice were housed in the specific pathogen free (SPF) animal facilities and fed with irradiated standard pellet chow and reverse osmosis water. All experiments were performed according to the French ethics committee regulations.

### Photothrombotic stroke model

Male mice were anesthetized by intraperitoneal injection of Ketamine/Xylazine and immobilized on a stereotaxic frame. An incision along the midline from the eye level down to the neck was made and the illuminated area on the exposed skull was determined corresponding to the following coordinates: 2.5mm posterior to bregma and 2.0mm lateral to midline. A fiber optic connected to KL 1600LED cold light source was placed on the targeting area after 100μl of Rose Bengal (10mg/ml) was injected. Illuminating for 15 minutes activated the Rose Bengal. After the skin was stitched, animals were put on a heating pad and returned to their home cages until they were fully awake.

### Beam-walk sensorimotor test

Mice were trained to walk across a 60cm long beam of 0.8 cm diameter, for a minimum of 4 days prior to surgery. For each trial, the mouse was placed at the start and allowed to cross the beam and was given 45 seconds to reach the goal cage between trials. Each mouse was given five consecutive trials on each test day. The total number of forelimb and hindlimb faults were scored by video recordings. For *in vivo* depletion of NK cells, mice were injected 100 μg of InVivoPlus anti-mouse Nk1.1 (Clone PK136, Bio X Cell) or InVivoPlus mouse IgG2a (Clone C1.18.4, Bio X Cell) resolved in PBS at P-9 and P-2, respectively.

### Administration of 4-hydroxytamoxifen

4-hydroxytamoxifen (4OHT - Sigma T176) was prepared in 667μl of ethanol and 6 ml of peanut oil. 160μl of 4OHT was administered intraperitoneally on 3 consecutive days(Jahn et al., 2018). Mice underwent 3 weeks of 4OHT before stroke induction.

### Flow cytometry

Brains were digested in HBSS containing 0.075mg/ml liberase TM and 0.2mg/ml Dnase l at 37℃ for 60 min. After being harvested and washed by HBSS containing 2% FBS, the brain cells suspended in 30% Percoll were centrifuged at 700 G for 15 min, resuspended in FACS buffer (HBSS containing 2% FBS). Dura maters isolated from skull as well as deep cervical, mandibular, axillary and mesenteric lymph nodes were digested in HBSS containing 0.075mg/ml liberase TM and 0.2mg/ml Dnase l at 37℃ for 20 min. Incubating collected blood with 1 X RBC lysis buffer (Invitrogen, 2139995) for 30s on ice, then stopping the reaction by HBSS containing 2% FBS. Cell suspensions were centrifuged at 300 G for 7 min and then resuspended in FACS buffer. In the FACS experiments for Fig. 2, S1 and S2, cells were blocked in FACS buffer containing 15% normal mouse serum for 15 min and subsequently stained for CD45 (BD biosciences, 564279), CD3 (BD biosciences, 562286), CD8 (BD biosciences, 562283), CD19 (BD biosciences, 562291), Ly6G (BD biosciences, 562700), F4/80 (BD biosciences, 565613), NKp46 (BD biosciences, 562850), CD127 (Ebioscience, 50-1271-80), ST2 (Ebioscience, 12-9333-80), KLRG1 (BD biosciences, 563595), CD4 (BD biosciences, 561099) diluted in FACS buffer for 30 min on ice. The live and dead cells were distinguished using LIVE/DEAD Fixable Blue Dead Cell Stain Kit (Life technologies, L23105). Cells were subsequently washed and resuspended in FACS buffer. Cells were blocked in FACS buffer containing 15% normal mouse serum for 15 min and subsequently stained for CD45, CD3e, CD8a, CD19, Ly6G, TCRβ, F4/80, NKp46, NK1.1, CD49a, CD49b, CD127 (Ebioscience, 50-1271-80), ST2, KLRG1 diluted in FACS buffer for 30 min on ice. Cells were washed with FACS buffer and resuspended in FACS buffer containing Sytox Blue. Samples were acquired on the BD LSR Fortessa-X20 cytometer and data were analyzed using FlowJo (version 10, LLC) software.

### Immunofluorescence

The mice were perfused with PBS containing 5U/ml of Heparin followed by 4% paraformaldehyde (PFA). The brains were extracted and fixed by 4% PFA overnight at 4℃. 50μm Vibratome sections were cut and blocked in EBT buffer (EBSS with 1% BSA and 2% Triton-X100) containing 10% serum for 1h at room temperature. Subsequently, sections were incubated with primary antibodies diluted in EBT buffer containing 3% serum for 24h at 4℃. After being washed 3 times with PBS containing 0.2% Triton-X100, the sections were incubated with fluorochrome-coupled secondary antibodies diluted in EBT buffer containing 3% serum for 12h at 4℃. After being washed 3 times with PBS, the sections were cleared in Histodenz medium (4g Histodenz (Sigma D2158) in 3 ml of 0.02M phosphate buffer) for 24h at room temperature and then mounted in Histodenz medium. Confocal images were acquired on a Zeiss-LSM880 confocal microscope and processed by ImageJ.

### Nissl staining

Vibratome sections were cut and mounted on positive charged plus slides (ThermoFisher scientific, SuperFrost Plus). The slides were placed in 1:1 alcohol/chloroform overnight and then rehydrated through 100% and 95% alcohol. Then the slides were placed into 0.1% cresyl violet for 10 minutes and rinsed quickly in distilled water. The sections were cleared with xylene for 10 minutes after being differentiated in 95% ethyl alcohol for 5 minutes and dehydrated in 100% alcohol for 10 minutes.

### Whole mount immunofluorescence

The *RORc^eGFP^* mice were perfused with PBS containing 5U/ml of Heparin followed by 0.4% paraformaldehyde (PFA). The brains were harvested and fixed by 0.4% PFA overnight at 4℃ and subsequently placed in permeabilization solution for 3 days. Subsequently, they were incubated in blocking medium PBS-MT (1% skim milk, 0.4% Triton X-100, 5% NDS and 5% NGS) for 3 days and subsequently incubated with anti-GFP (AVES, GFP-1020), anti-CD3 (BD biosciences, 563024), anti-KLRG1 (BD biosciences, 563595) and anti-NKp46 (BD biosciences, 562850) in PBS-MT with 3% NDS plus NGS for 10 days. After being washed with PBS containing 0.2% Triton X-100, brains were incubated with donkey anti-chicken Cy3 (Jackson Immunoresearch, 703-166-155), goat anti-armenian hamster 647 (Jackson Immunoresearch, 127-605-160), goat anti-Syrian hamster 488 and donkey anti-rat 790 (Jackson Immunoresearch, 712-655-153) in PBS-MT with 3% NDS as well as NGS for 10 days. Brains were washed with PBS and dehydrated by methanol series (20%, 40%, 60%, 80% 1h each and 100% 2h). Subsequently, the brains were placed in 66% dicholoromethane (Sigma, 270997), 33% methanol overnight, and afterwards in 100% dicholoromethane for 15 min. The brains were cleared in BABB (2:1 benzyl alcohol (Honeywell, 108006)/benzyl benzoate (Acros organics, 105862500)) overnight. Brains were acquired using the light sheet Ultramicroscope version II (LaVision BioTec). 3D imaging analysis were performed using Imaris software (Version 9.1.0, Bitplane).

### Quantitative PCR

Total RNA was prepared by TRIzol reagent (Sigma). cDNA synthesis was carried out using RevertAid RT Kit (ThermoFisher scientific). The transcripts were amplified on the 7500 Fast Real-Time PCR System. The amplification of *Cxcr4* and the corresponding *Gapdh* was detected by Taqman probes (Cxcr4-Mm01292123_m1, Gapdh-Mm99999915_g1; ThermoFisher scientific), combining with Taqman Fast Advanced Master Mix (ThermoFisher scientific). The amplification of *Cxcl12* and the corresponding *Gapdh* was detected by the following primers: *Cxcl12*_forward 5’ - tcaagcatctgaaaatcctcaaca-3’, *Cxcl12*_reverse 5’-ttcgggtcaatgcacacttgt-3’, *Gapdh*_forward 5’-tgtgtccgtcgtggatctga-3’, *Gapdh*_reverse 5’-cctgcttcaccaccttcttga-3’, combining with TB Green Premix Ex Taq (Takara). Relative expression was calculated by ΔΔC_T_ method.

### Statistical analysis

Graphs and statistical significance of differences between groups by two-way ANOVA in all figures were analyzed using GraphPad Prism (v.7.03) software. All the cartoons were generated using BioRender.

### Staining panels for FACS

Staining panel I for FACS in Fig. 2, S1 and S2

|  |  |  |
| --- | --- | --- |
| CD45 | BV395 | BD biosciences, 564279 |
| CD3 | PE-CF594 | BD biosciences, 562286 |
| CD8 | PE-CF594 | BD biosciences, 562283 |
| CD19 | PE-CF594 | BD biosciences, 562291 |
| Ly6G | PE-CF594 | BD biosciences, 562700 |
| F4/80 | PE-CF594 | BD biosciences, 565613 |
| NKp46 | BV421 | BD biosciences, 562850 |
| CD127 | eFluor 660 | Ebioscience, 50 – 1271 – 80 |
| ST2 | PE | Ebioscience, 12 – 9333 – 80 |
| KLRG1 | PerCP-Cy5.5 | BD biosciences, 563595 |
| CD4 | PE-Cy7 | BD biosciences, 561099 |
| RORyt | RORyt-eGFP | Sparwasser et al., 2004 |
| ReporterLive/Dead | Fixable Blue Dead Cell Stain Kit | Life technologies, L23105 |

Staining panel II for FACS in Fig. 3, 4, 5, S4, S5, S7, S8 and S9

Staining panel II for FACS in Fig. 3, 4, 5, S4, S5, S7, S8 and S9
|  |  |  |
| --- | --- | --- |
| CD45 | BV395 | BD biosciences, 564279 |
| CD3 | BV510 | BD biosciences, 563024 |
| CD8 | V500 | BD biosciences, 560776 |
| CD19 | BV510 | BD biosciences, 562956 |
| Ly6G | BV510 | Biolegend, BLE127633 |
| F4/80 | BV510 | BD biosciences, 743280 |
| TCRβ | BV510 | BD biosciences, 563221 |
| NKp46 | BV421 | BD biosciences, 562850 |
| NK1.1 | Alexa Fluor 700 | Ebioscience, 56 – 5971 – 82 |
| CD49a | BV711 | BD biosciences, 564863 |
| CD49b | PE-Cy7 | Miltenyi, 130 – 116 – 370 |
| CD127 | eFluor 660 | Ebioscience, 50 – 1271 – 80 |
| ST2 | PE | Ebioscience, 12 – 9333 – 80 |
| KLRG1 | PerCP-Cy5.5 | BD biosciences, 563595 |
| Live/Dead | Sytox Blue | Life technologies, S34857 |

## Acknowledgements

The work was supported by the FRM Amorçage jeunes équipes (AJE20150633331), ANR ACHN (ANR-16-ACHN-0011), ANR PRCI (ANR-17-CE13-0029-01), A*midex Chaire d’excellence to SvdP and institutional grants to the CIML from INSERM, CNRS and Aix-Marseille University. We thank the animal facility, notably Toufik Guelmami and Michel Pontier. The flow cytometry core facility, notably Marc Barad, Sylvain Bigot and Laurence Borge. We acknowledge the PICSL imaging facility of the CIML (ImagImm), member of the national infrastructure France-BioImaging supported by the French National Research Agency (ANR-10-INBS-04), notably Mathieu Fallet and Sebastian Mailfert, Lionel Chasson of the histology platform. Gérard Eberl (Pasteur Institute, Paris, France) for kindly supplying the *RORc^eGFP^* mice, Ralf Adams (Max Planck Institute for Molecular Medicine, Münster, Germany) for the *Cdh5^CreERT2^* mice and Eric Vivier (CIML, Marseille, France) for the *Ncr1^iCre^* mice.

The authors declare no conflict of interest.

