## Supplemental Figures for "Brain endothelial CXCL12 attracts protective Natural Killer cells during ischemic stroke"

Supplementary Figure Legends

**Supplementary Figure 1: Gating strategy for flow cytometry analysis on *RORc^eGFP^*-reporter mice.** Gating strategy for FACS on brain hemisphere with the lesion (A), dura mater (B) and lymph node (C) samples, NK ($CD45^+Lin^-NKp46^+CD127^-$), ILC1 ($CD45^+Lin^-NKp46^+CD127^+ROR\gamma t^-$), ILC2 ($CD45^+Lin^-NKp46^-CD127^+ROR\gamma t^-ST2^+KLRG1^+$), $NKp46^+$ILC3 ($CD45^+Lin^-NKp46^+CD127^+ROR\gamma t^+$), LTi-like ($CD45^+Lin^-NKp46^-CD127^+ROR\gamma t^+$), and Th17 ($CD45^+Lin^+CD4^+ROR\gamma t^+$). Representative dot plots of ILC and Th17 cells at day 15 are shown.

**Supplementary Figure 2: Quantification of ILCs and Th17 cells in the dura mater and lymph nodes after PT induction.** FACS analysis of ILCs in the dura mater (A), NK ($CD45^+Lin^-NKp46^+CD127^-$), ILC1 ($CD45^+Lin^-NKp46^+CD127^+ROR\gamma t^-$), ILC2 ($CD45^+Lin^-NKp46^-CD127^+ROR\gamma t^-ST2^+KLRG1^+$), $NKp46^+$ILC3 ($CD45^+Lin^-NKp46^+CD127^+ROR\gamma t^+$), LTi-like ($CD45^+Lin^-NKp46^-CD127^+ROR\gamma t^+$), and Th17 ($CD45^+Lin^+CD4^+ROR\gamma t^+$). Representative dot plots of ILCs and Th17 cells at P15 are shown. (B) Quantification of ILC populations and Th17 cells in the dura mater at different time points after PT induction. Each data point represents an individual mouse. (C) Flow cytometry analysis of ILCs in the lymph nodes, representative dot plots of ILCs and Th17 cells at day 15 are shown. Quantification of ILC and Th17 cells in the lymph nodes including (D) deep cervical lymph node (dcLN), (E) mandibular LN (mandiLN), (F) mesenteric LN (mLN) and (G) axillary LN (axiLN). Each data point represents an individual mouse.

**Supplementary Figure 3: Sorting endothelial cells.** Gating strategy for sorting endothelial cells ($CD31^+CD45^-$) from WT (*Cxcl12$^{fl/fl}$*) (A) and KO (*Cxcl12$^{Cdh5-/-}$*) (B) mice.

**Supplementary Figure 4: Gating strategy for flow cytometry on *Cdh5$^{CreERT2}$;Cxcl12$^{fl}$* and *Ncr1$^{iCre}$;Cxcr4$^{fl}$* mice.** Gating strategy for FACS on brain hemisphere with the lesion (A), dura mater (B) and lymph node (C) samples; NK ($CD45^+Lin^-NKp46^+NK1.1^+CD49a^-CD49b^+$), intILC1 ($CD45^+Lin^-NKp46^+NK1.1^+CD49a^+CD49b^+$), ILC1

($CD45^+Lin^-NKp46^+NK1.1^+CD49a^+CD49b^-$), ILC2 ($CD45^+Lin^-NKp46^-NK1.1^-CD127^+ST2^+KLRG1^+$) and $NKp46^+$ILC3 ($CD45^+Lin^-NK1.1^-NKp46^+CD127^+$). Representative dot plots of ILCs at day 15 are shown.

**Supplementary Figure 5: Quantification of ILCs in the dura mater and lymph nodes from *Cxcx12*-deficient mice after PT.** (A) Flow cytometry analysis of ILCs in the dura mater from *Cxcl12*$^{fl/fl}$ and *Cxcl12*$^{Cdh5–/–}$ mice at day 15 after PT induction. Representative dot plots of ILCs at P15 are shown. (B) Quantification of ILCs in the dura mater at P2 and P15. Each data point represents an individual mouse. NK *P=0.0483 (C) Flow cytometry analysis of ILCs in the lymph node from *Cxcl12*$^{fl/fl}$ and *Cxcl12*$^{Cdh5–/–}$ mice at P15. Representative dot plots of ILCs at P15 are shown. Quantification of ILCs in the lymph nodes including (D) deep cervical lymph node (dcLN), (E) mandibular LN (mandiLN), (F) mesenteric LN (mLN) and (G) axillary LN (axiLN). Each data point represents an individual mouse. Statistics: (E) NK *P=0.105; ILC1 **P=0.0043; (G) $NKp46^+$ ILC3 *P=0.0344.

**Supplementary Figure 6: Sorting $NKp46^+$ ILC cells.** Gating strategy for sorting $NKp46^+$ ILC cells ($CD45^+Lin^-NKp46^+$) from WT (*Cxcr4*$^{fl/fl}$) (A) and KO (*Cxcr4*$^{Ncr1-/-}$) (B) mice. Lin: CD3e, CD8a, CD19, Ly6G, TCRβ, F4/80.

**Supplementary Figure 7: Analysis of $NK1.1^+NKp46^+$ ILC cell numbers in the *Cxcr4*$^{fl/fl}$ and *Ncr*$^{iCre/+}$ mice**. (A) FACS analysis of NK and ILC1 cells in the liver of *Cxcr4*$^{fl/fl}$ and *Ncr*$^{iCre/+}$ mice. (B) Analysis of NK cells in the PT stroke brain of *Cxcr4*$^{fl/fl}$ and *Ncr*$^{iCre/+}$ mice, at P0, P2 and P15. Each data point represents an individual mouse.

**Supplementary Figure 8: Quantification of ILCs in the dura mater and lymph nodes in *Cxcr4*$^{Ncr1-/-}$ mice.** (A) FACS analysis of ILCs in the dura mater from *Cxcr4*$^{fl/fl}$ and *Cxcr4*$^{Ncr1-/-}$ mice at P15. Representative dot plots of ILCs at P15 are shown. (B) Quantification of ILCs in the dura mater at P0, P2 and P15. Each data point represents an individual mouse. NK P2 *P=0.0107, P15 ***P=0.0004; ILC1 P15 *P=0.0419. (C) Flow cytometry analysis of ILCs in the lymph nodes from *Cxcr4*$^{fl/fl}$ and *Cxcr4*$^{Ncr1-/-}$ mice at P15. Representative dot plots of ILCs at P15 are shown. Quantification of ILCs in the lymph nodes including (D) deep cervical lymph node (dcLN), (E) mandibular LN (mandiLN), (F) mesenteric LN (mLN) and (G) axillary LN (axiLN). Each data point represents an individual mouse. Statistics: (D) NK P2 **P=0.0078; (F) NK P0 *P=0.0137, P2 ***P=0.0003, P15 *P=0.0141.

**Supplementary Figure 9: Gating strategy for analyzing the presence of NK cells in the blood.** Lin: CD3e, CD8a, CD19, Ly6G, TCRβ, F4/80.

**Supplementary video 1：3D imaging for brain lesion from stroke mouse.** Whole mount staining for the stroke brain at day 2 after PT induction. In the original acquisition KLRG1 in shown in cyan, RORγt in green, CD3 in red, NKp46 in white. Using the spot function in Imaris, we determined NK cells, ILC1s and $NKp46^{+}$ILC3s ($CD3^{-}$ $NKp46^{+}$) as white spots, ILC3s and LTi cells ($CD3^{-}$$ROR\gamma t^{+}$) as green spots and ILC2s ($CD3^{-}$ $KLRG1^{+}$) as cyan spots.

**Supplementary video 2：3D imaging for brain lesion from stroke mouse.** Whole mount staining for the stroke brain at day 10 after PT induction. In the original acquisition KLRG1 in shown in cyan, RORγt in green, CD3 in red, NKp46 in white. Using the spot function in Imaris, we determined NK cells, ILC1s and $NKp46^{+}$ILC3s ($CD3^{-}$ $NKp46^{+}$) as white spots, ILC3s and LTi cells ($CD3^{-}$$ROR\gamma t^{+}$) as green spots and ILC2s ($CD3^{-}$ $KLRG1^{+}$) as cyan spots.

### Supplementary Figure 1

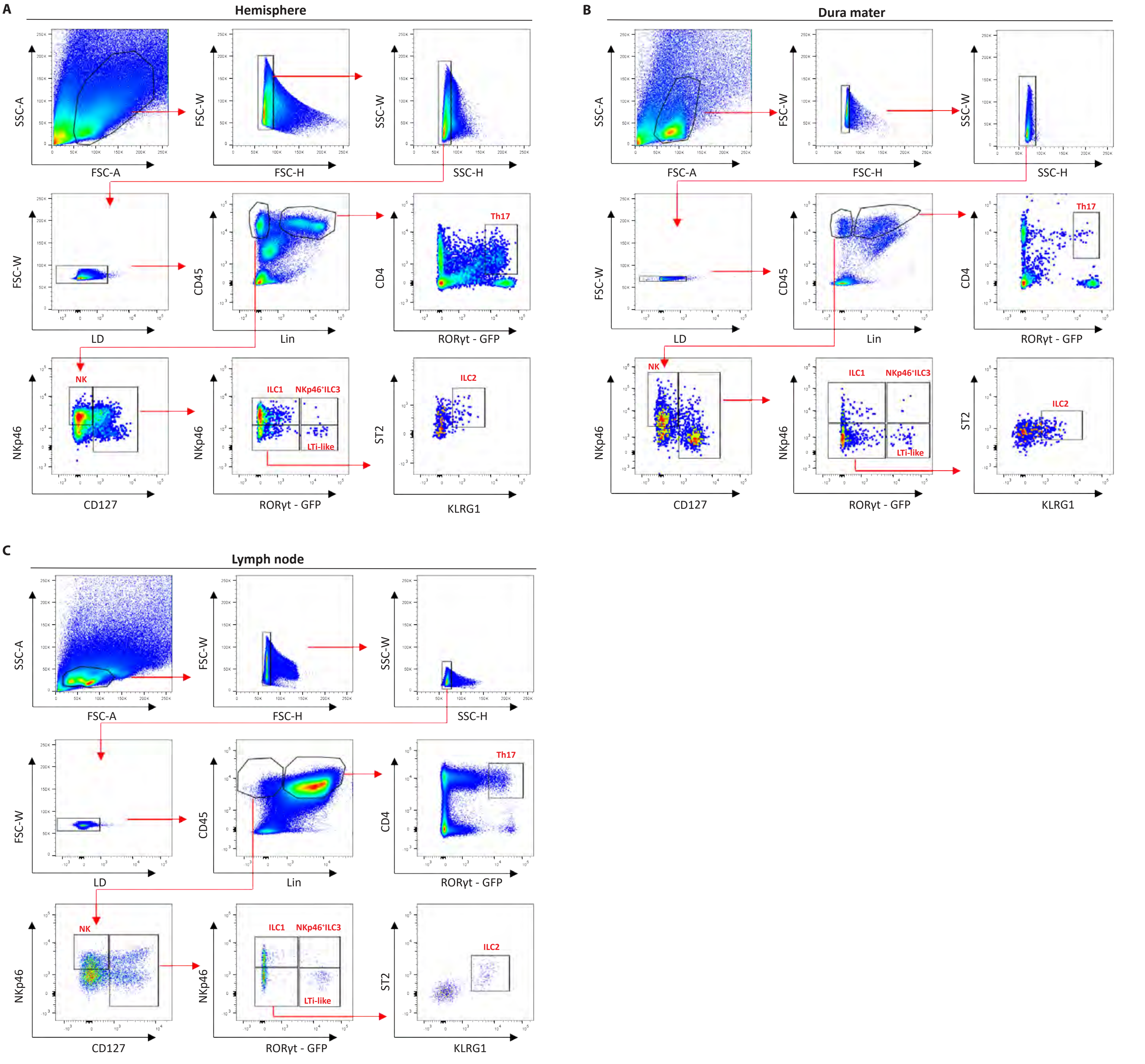

### Supplementary Figure 2

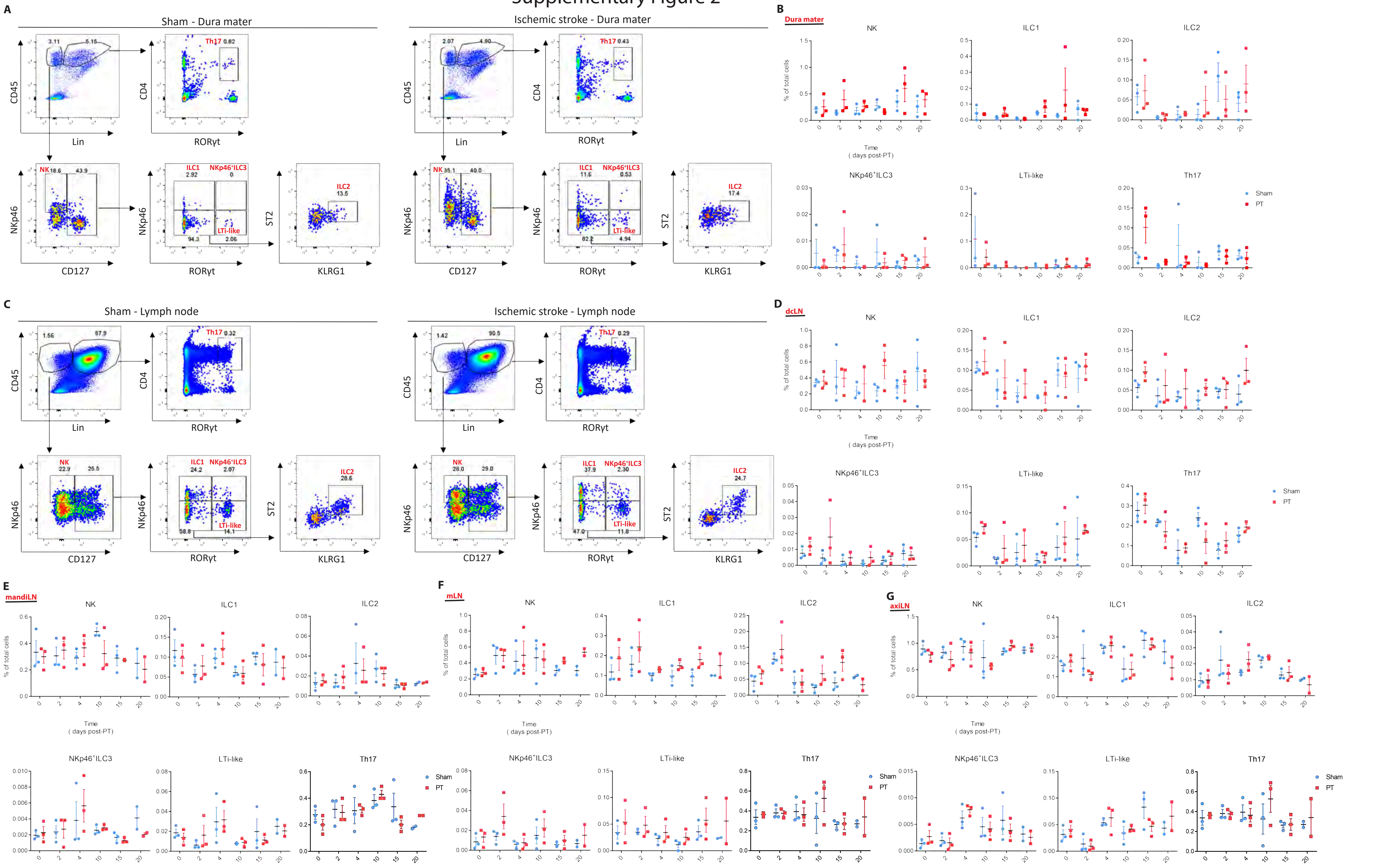

### Supplementary Figure 3

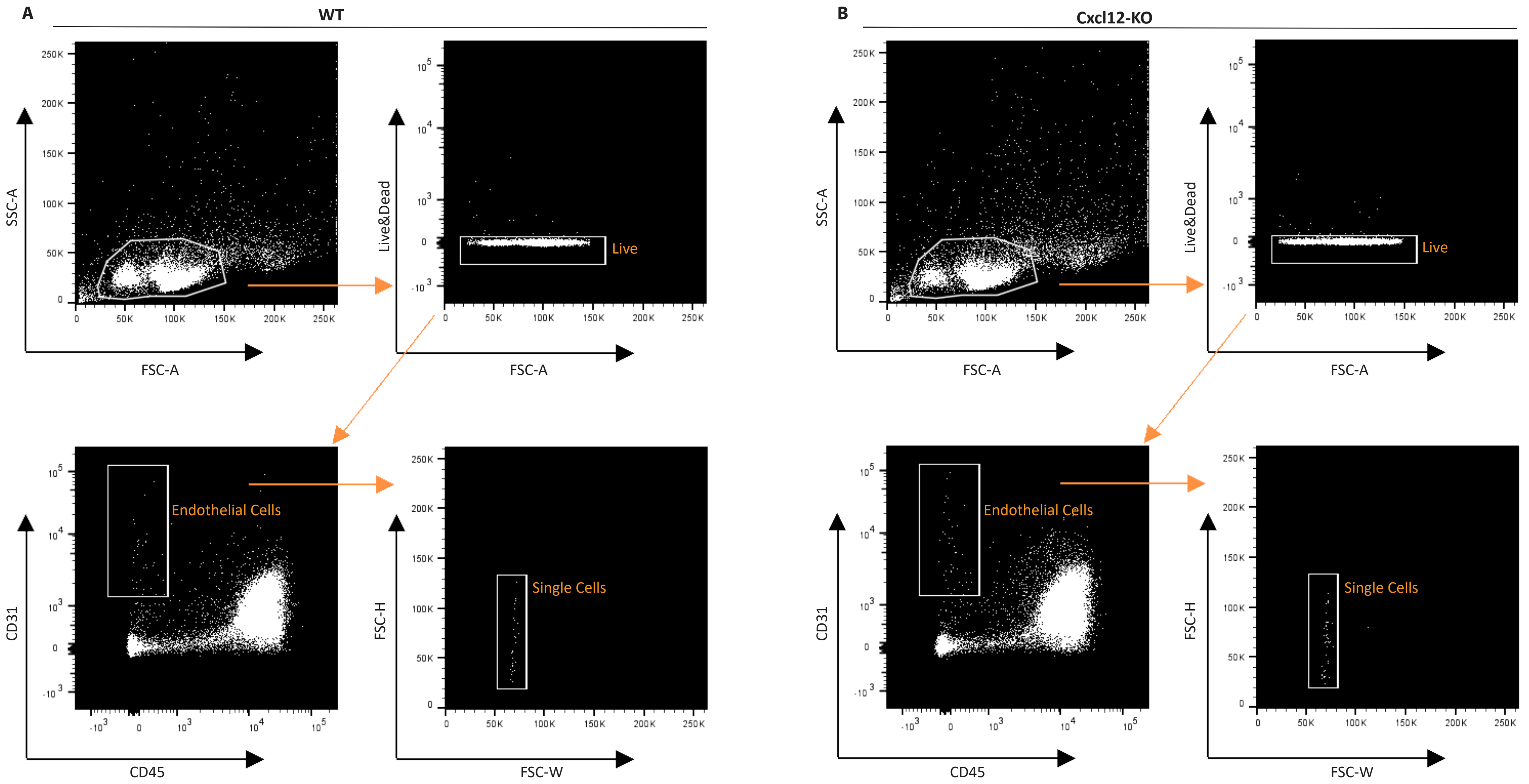

### Supplementary Figure 4

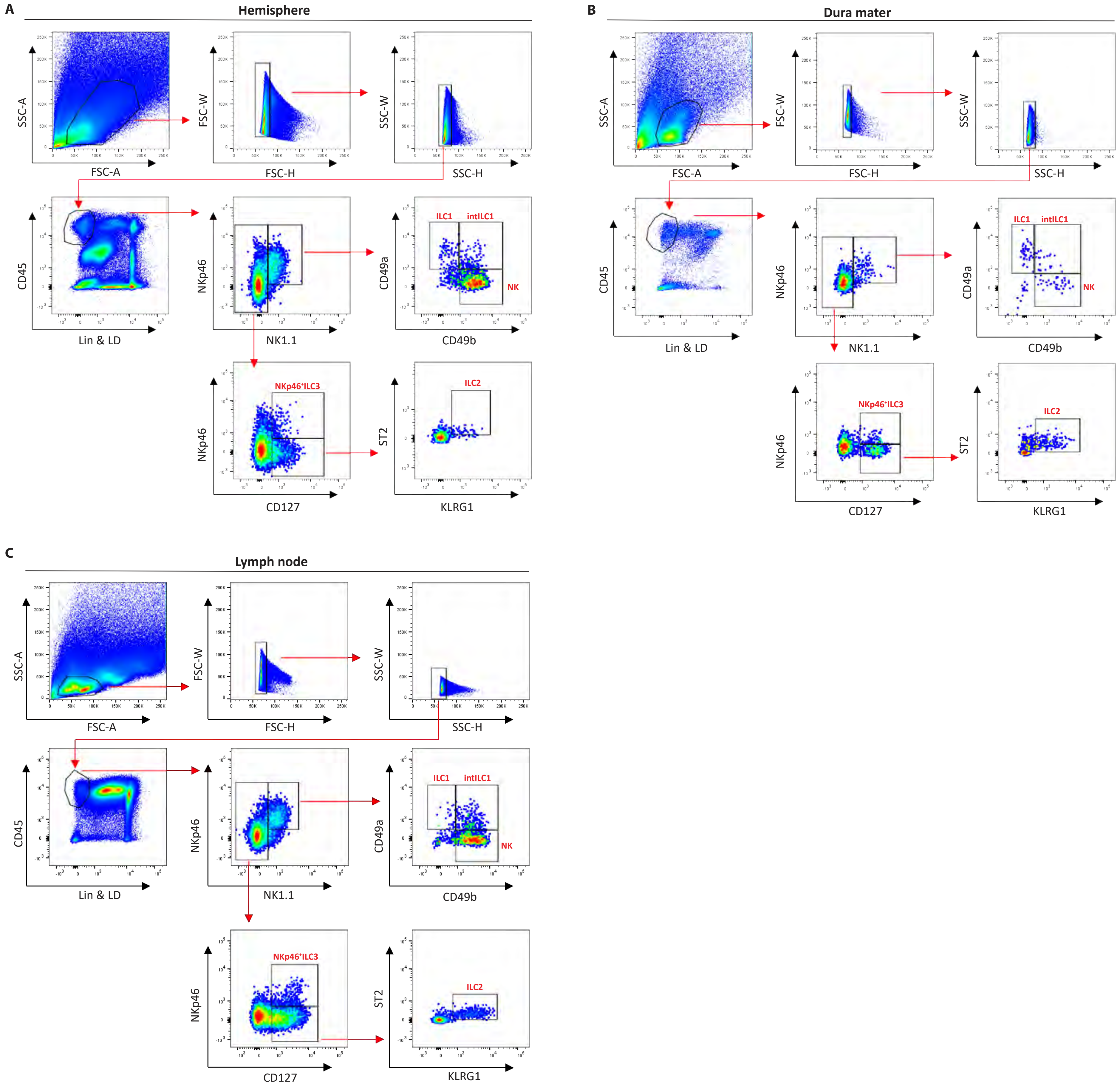

### Supplementary Figure 5

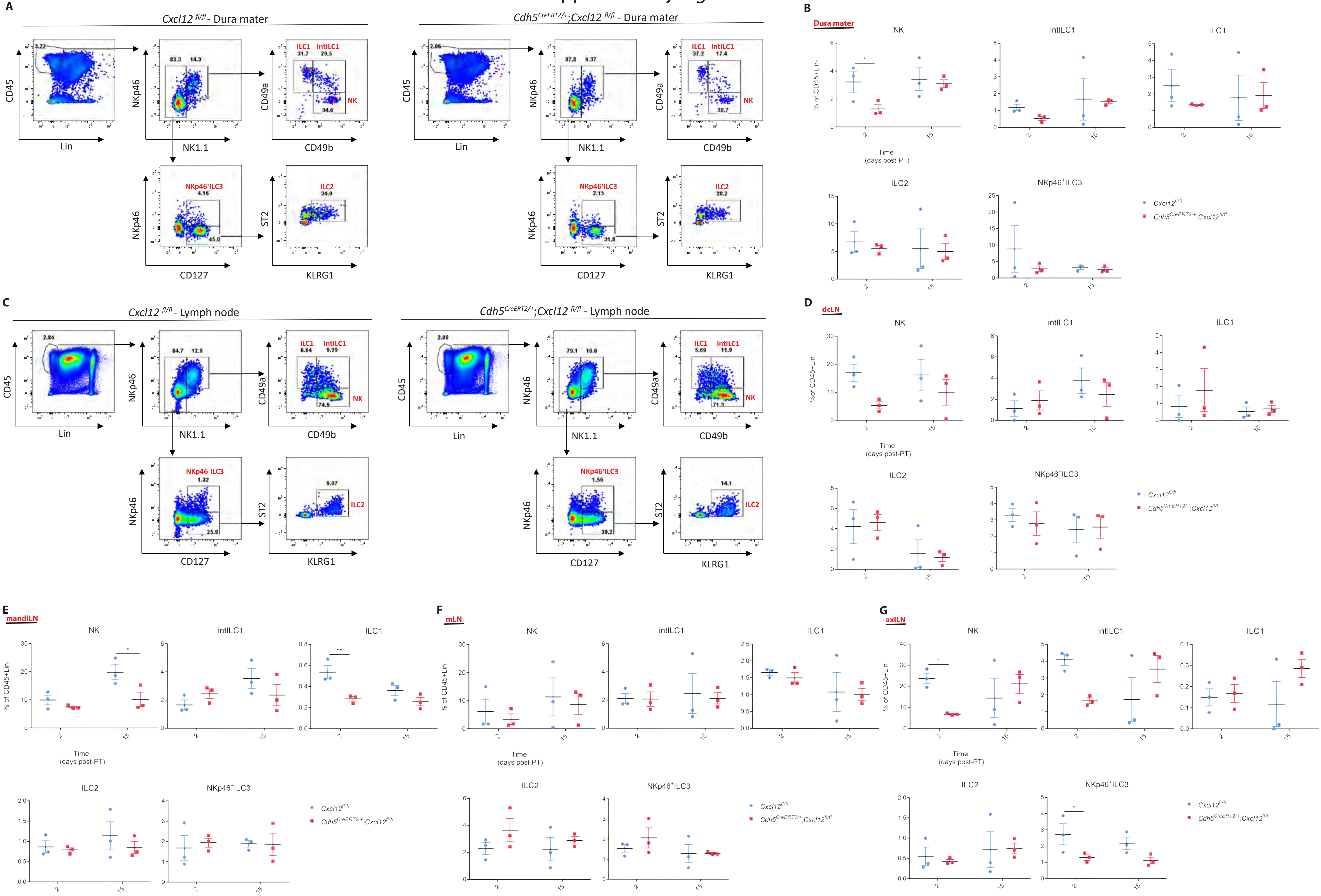

### Supplementary Figure 6

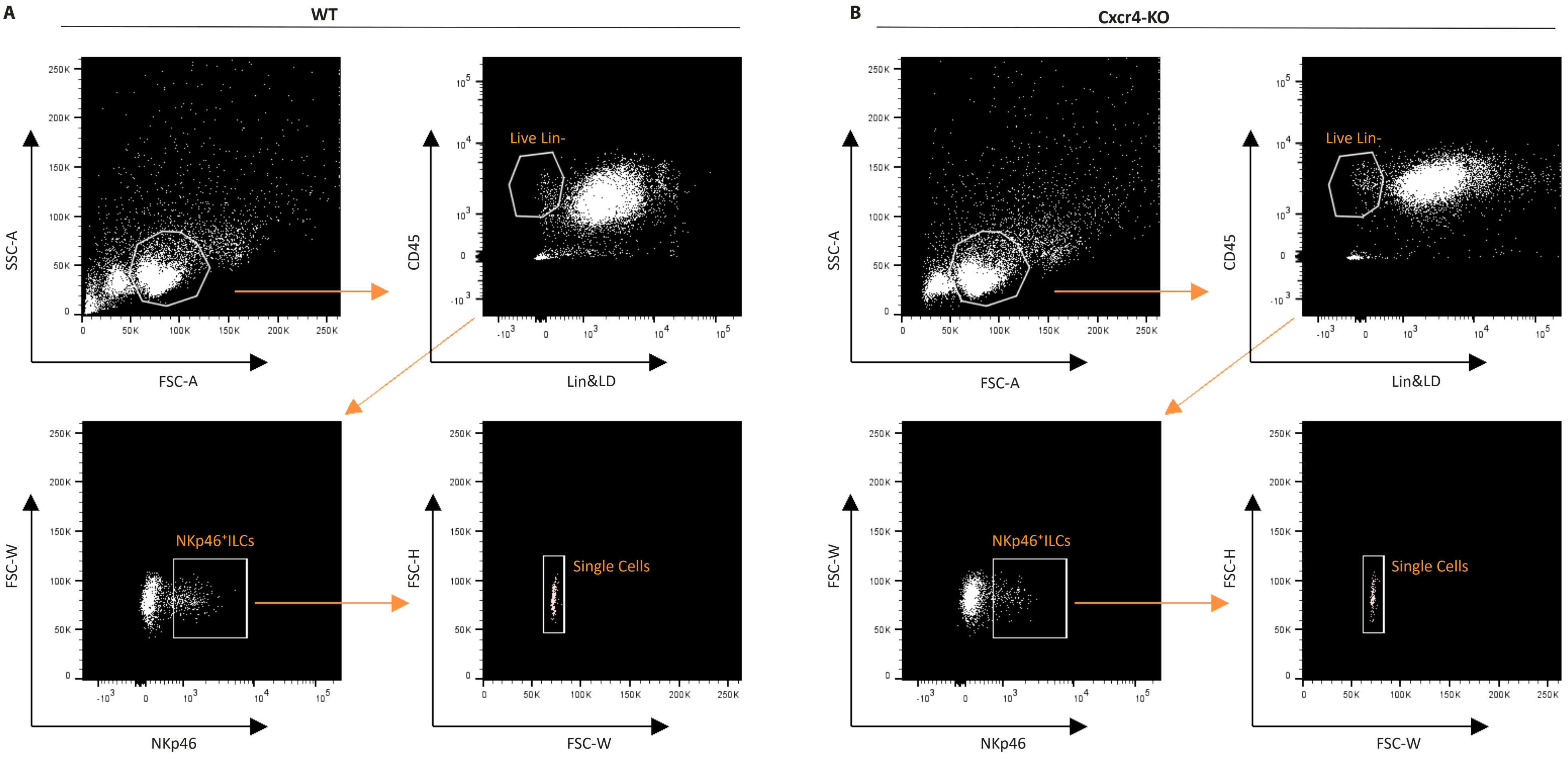

### Supplementary Figure 7

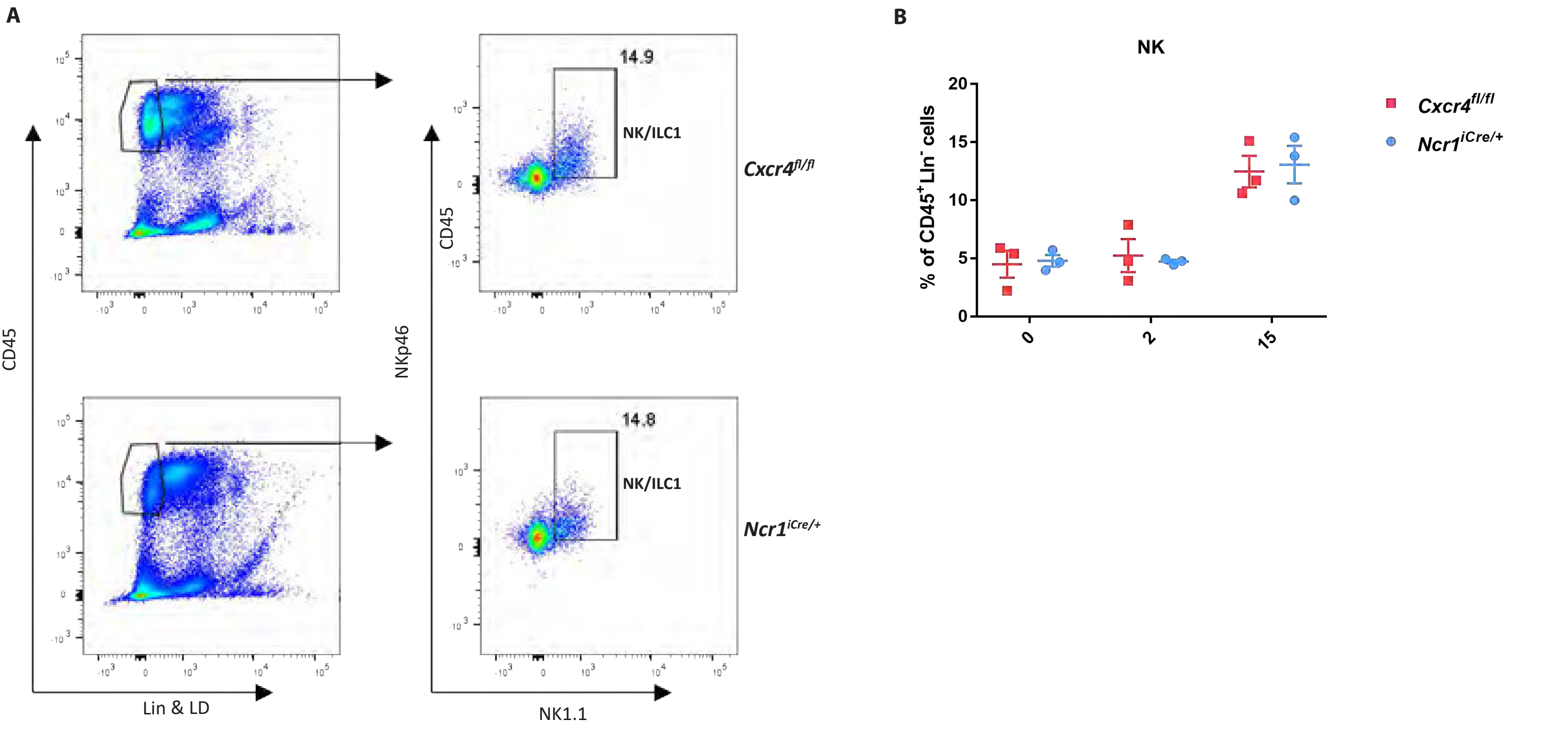

### Supplementary Figure 8

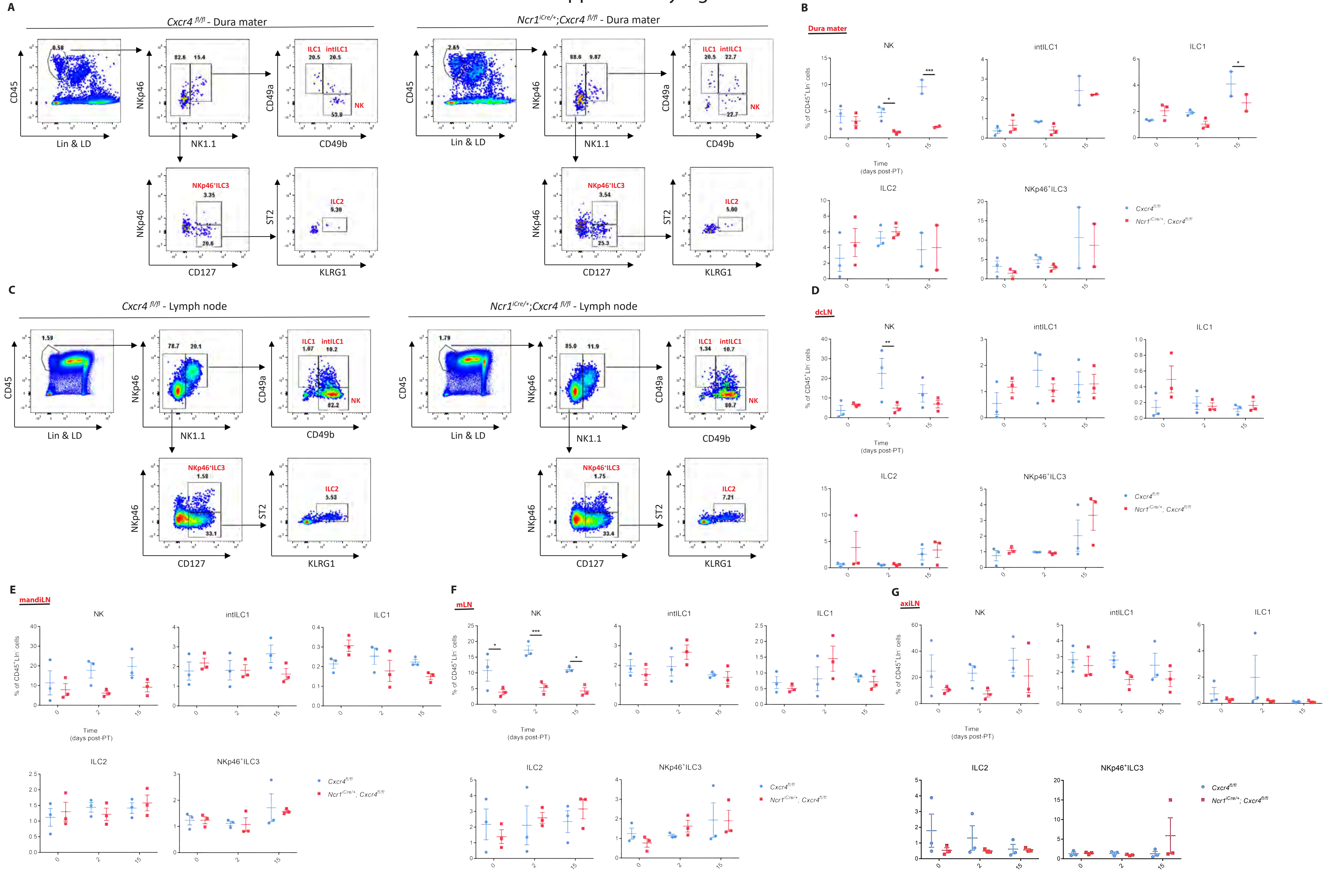

### Supplementary Figure 9

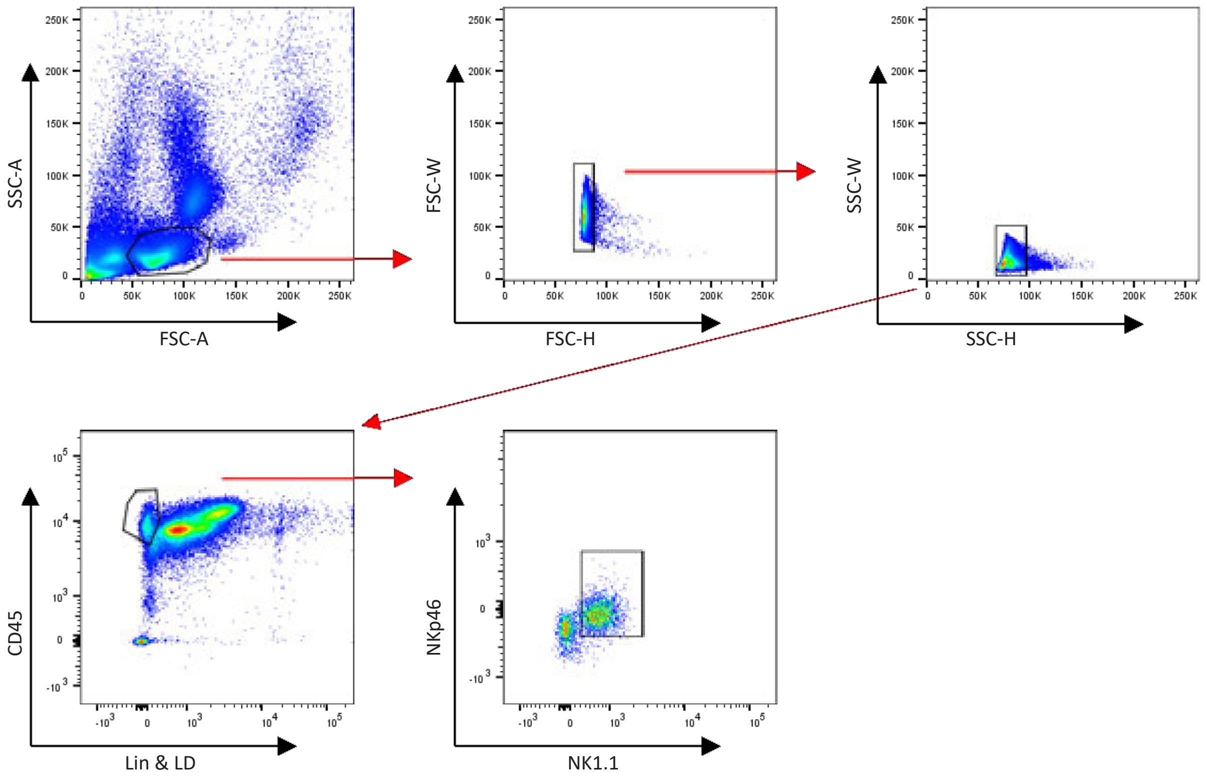
